# TrES-seq integrates single-cell RNA and nanobody-based histone mark profiles to decode epigenetic and transcriptional axes of cell identity

**DOI:** 10.64898/2026.11.20.767314

**Authors:** Vamsi Krishna Polavarapu, Ahrmad Annan, Divyanshu Srivastava, Natalya Katanayeva, Justine Epiney, Albert Santamaria-Martinez, Karma Lambercy, Giovanni Ciriello, Elisa Oricchio

**Affiliations:** Swiss Institute for Experimental Cancer Research (ISREC), School of Life Sciences, EPFL, Lausanne, Switzerland; Swiss Cancer Center Leman (SCCL), Lausanne, Switzerland; Department of Computational Biology, University of Lausanne, Lausanne, Switzerland; Swiss Institute of Bioinformatics, Lausanne, Switzerland; Lausanne University Hospital (CHUV), Switzerland

## Abstract

Histone post-translational modifications and gene expression jointly shape cellular identity and activity. However, systematically assessing their heterogeneity and functional interdependence requires high-quality multimodal single-cell readouts, whose generation still faces significant technological challenges. Here, we developed TrES-seq (transcriptional and epigenetic state sequencing), a method that integrates scRNA-seq with species-specific nanobody–Tn5 fusion proteins to jointly profile RNA and two histone modifications in the same cells and across multiple samples. In human tonsils, single-cell H3K27ac, H3K27me3, H3K9me3, and RNA data in >25,000 cells revealed a “one-to-many” relationship where heterogeneous transcriptional states shared similar epigenetic profiles. For example, naïve and memory B cells were transcriptionally different but epigenetically indistinguishable, while the epigenetic status of enhancers but not promoters separated germinal center from naïve and memory B cells. Moreover, naturally occurring and treatment-induced cell state transitions revealed broad transcriptional changes occurring among genes that already exhibited and retained an epigenetically active state. Overall, TrES-seq provides a robust and cost-effective platform to determine how epigenetic states influence transcriptional heterogeneity, shaping cell phenotypes and function in different physiological and pathological contexts.

## Introduction

During cell differentiation, transcriptional and epigenetic changes guide the development of hundreds of distinct cell types from pluripotent stem cells^1,2^. Immortalized in the valleys and peaks of Waddington’s landscape, distinct lineage trajectories are the synthesis of the dynamic regulation of gene expression^3^. Indeed, large-scale single-cell transcriptomic profiling has revealed a wide variety of cell types and subtypes^4,5^, and dynamic transitions among them occurring in response to different stimuli and conditions^6,7^. These transitions can be reversible, including de-differentiation and trans-differentiation pathways, and exist along a continuous spectrum of cell states^8,9^. To what extent diverse epigenetic states mirror such transcriptional heterogeneity has been more challenging to determine. The integration of independent experiments measuring either single-cell RNA, chromatin accessibility, or individual histone marks on matching samples provided the first insight into the functional interplay between these molecular layers^10^. For example, changes in open and closed chromatin or between repressing and activating histone marks were suggested to anticipate transcriptional reprogramming^11,12^. However, these approaches could only infer a temporal association between epigenetic features and gene expression, lacking concurrent measurements in the same cells.

To fill this gap, pioneering multimodal approaches were developed to measure RNA abundance and chromatin accessibility via Assay for Transposase-Accessible Chromatin (scRNA+ATAC-seq) in the same cells^13,14^. These technologies revealed asynchronous changes between epigenetic and transcriptional states, demonstrating, for example, that chromatin accessibility changes preceded and possibly allowed to predict transcriptional activation^15^. Nevertheless, chromatin accessibility assays oversimplify the epigenome into a combination of “open” and “closed” states. In contrast, histone post-translational modifications (HPTMs), either alone or in combination, allow the definition of a more granular spectrum of chromatin states associated with distinct regulatory functions^16^. Although more informative, single-cell HPTM profiles remain scarce because current technologies face significant challenges. For example, technologies such as scCUT&Tag^17^, nano-CUT&Tag^18^ (nano-CT), and NTT-seq^19^ use species-specific antibodies or nanobodies to assay one or two histone marks in the same cell. However, these methods cannot measure RNA abundance and adopt expensive microfluidic droplet strategies to barcode individual cells. More recently, a few multimodal approaches have been proposed to combine HPTM profiling with scRNA-seq. These include PAIRED-Tag^20^ and OneCell CUT&Tag^21^, which measure one histone mark and gene expression and revealed different transition dynamics between epigenetic and transcriptional states. uCoTargetX^22^ and scMTR-seq^23^ extended these approaches to multiple HPTMs. However, to increase the number of histone marks detected, these approaches pre-mix antibodies with the tagging enzyme, which might reduce specificity.

To overcome these challenges, we developed a new method that simultaneously assays RNA and two HPTMs in single cells to define their **Tr**anscriptional and **E**pigenetic **S**tate by sequencing (**TrES**-seq). TrES-seq adopts species-specific nanobody-Tn5 fusion proteins to minimize cross-contamination, and a split-and-pool barcoding strategy to reduce costs compared to microfluidic devices. We optimized TrES-seq to generate high-quality multimodal data for diverse combinations of HPTMs in multiple samples and conditions. Interestingly, multimodal profiles of primary human tonsils revealed distinct associations between epigenetic and transcriptional features of cell types, subtypes, and transitional cell states.

## Results

### Combined single-cell dual nano-CUT&Tag and gene expression

To assess the performance of multimodal technologies measuring at least two histone marks and gene expression, we implemented scMTR-seq and uCoTargetX to measure H3K27ac, H3K27me3, and RNA in the human germinal center B-cell lymphoma cell line, Karpas-422. Both scMTR-seq and uCoTargetX pre-assemble proteinA-Tn5 (pA-Tn5) antibodies and use a split-and-pool strategy, but they differ in their approach to reduce cross-contamination: MTR-seq applies all antibodies at the same time and uses IgG to block free pA-Tn5, while uCoTargetX applies pA-Tn5 complexes one at a time, washing unbound complexes between cycles (**Fig. 1a**). We tested both protocols to profile H3K27ac, H3K27me3, and RNA using either a short (1h) or a long (12h) incubation with pre-assembled pA-Tn5 and antibody complexes, although in the original protocol only 1h incubation was indicated. For scMTR-seq, H3K27ac data quality was poor with both incubation times, while we obtained good H3K27me3 profiles with 1h incubation (**Suppl. Figure 1a**). Conversely, with uCoTargetX, we obtained good data quality for both histone marks only after 12h incubation (**Suppl. Figure 1b**). Nevertheless, we still noticed substantial cross-contamination between H3K27ac and H3K27me3 signals with both technologies (**Fig. 1b**). Upon re-analyzing publicly available data generated with scMTR-seq in human pluripotent stem cells (hPSCs) and processed signal tracks for uCoTargetX (as the processed single-cell data or the code to demultiplex the original raw data was not made available), we confirmed extensive cross-contamination in both markers (**Suppl. Figure 1c**). In contrast to these protocols, to reduce cross-contamination and improve signal specificity, nano-CT relies on Tn5 species-specific nanobodies (**Fig. 1c**). Hence, we used the nano-CT protocol, and in both newly generated Karpas-422 profiles and the original publicly available data, cross-contamination was significantly reduced, although not eliminated in the H3K27me3 modality. (**Fig. 1d** and **Suppl. Figure 1d**). However, nano-CT lacks RNA information, and it relies on the 10X Genomics microfluidic kit to barcode individual cells.

**Figure 1.**
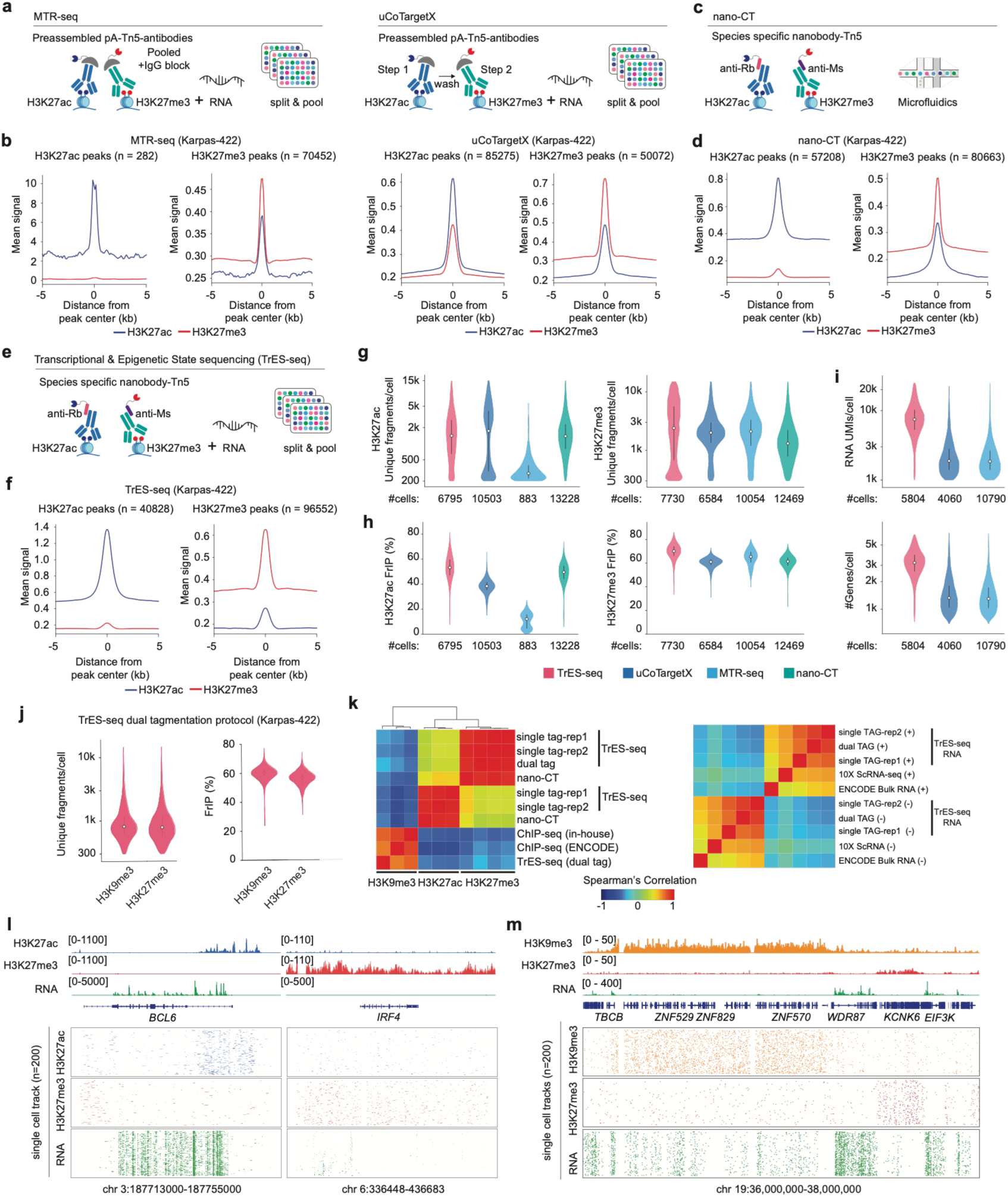
Benchmarking TrES-seq. **a.** Schematic representation of the scMTR-seq and uCoTargetX protocol. **b**. Mean H3K27ac and H3K27me3 coverage signal within ±5 kb of peak midpoints for MTR-seq and uCoTargetX. Profiles are centered on each dataset’s own peaks for the indicated mark. n= number of peaks. **c**. Schematic representation of the nano-CT (Nano CUT&Tag). **d**. Mean H3K27ac and H3K27me3 coverage signal within ±5 kb of peak midpoints for nano-CT. Profiles are centered on the peaks for the indicated mark. n= number of peaks. **e**. Schematic representation of TrES-seq. **f.** Mean H3K27ac and H3K27me3 coverage signal within ±5 kb of peak midpoints for TrES-seq. Profiles are centered on the peaks for the indicated mark. n= number of peaks. **g-i** Distributions of single-cell data quality metrics across the indicated technologies in Karpas-422 cells. (g) Unique H3K27ac (left) and H3K27me3 (right) fragments per cell. (h) Fraction of reads in H3K27ac peaks (left) and H3K27me3 peaks (right). (i) UMIs (top) and detected genes (bottom) per cell. **j**. Distributions of unique fragments per cell and fraction of reads in peaks for H3K27me3 and H3K9me3 in Karpas-422 cells profiled by dual tagmentation. Violin plots showing unique fragments per cell (left) and fraction of reads in peaks for both H3K27me3 and H3K9me3 produced using dual tagmentation protocol in Karpas-422. **k**. Left, heatmap representation of the Spearman’s correlations between whole-genome profiles of H3K27ac, H3K27me3, and H3K9me3 (column labels) generated using single-TAG or dual-TAG TrES-seq or independent technologies (row labels) in Karpas-422 cells. Right, Spearman correlations between strand-specific RNA profiles in Karpas-422 cells generated using single-TAG or dual-TAG TrES-seq, 10x scRNA-seq or ENCODE bulk RNA-seq. The + and − symbols denote genomic strands. **l,m,** Genome browser views of matched TrES-seq profiles at selected loci in Karpas-422 cells. Top, aggregated signal tracks for H3K27me3 (red) and RNA (green) alongside H3K27ac (blue) in l or H3K9me3 (orange) in m, with gene annotations. Bottom, corresponding single-cell read tracks across the same loci, ordered by descending yield.

To limit cross-contamination, obtain both epigenetic and transcriptional data, and allow high sample multiplexing, we developed the TrES-seq protocol, which integrates a modified version of single-cell nano-CT with RNA sequencing and uses a combinatorial barcoding strategy to assign reads to individual cells. Briefly, we used species-specific nanobody-Tn5 fusion proteins to cut and tag genomic sequences marked by selected HPTMs with an adapter containing modality and sample barcodes. Then we labeled RNA during reverse transcription using a biotinylated primer carrying a matching sample barcode (**Figure 1e**). In total, TrES-seq uses up to 12 sample barcodes to allow profiling different samples in the same experiment. To distinguish individual cells, we used a library of 48 barcodes, which were randomly combined through three consecutive ligations (**Suppl. Figure 1e** and **Suppl. Table 1**). The RNA was separated from DNA via biotin pulldown, and libraries were prepared for sequencing (**Suppl. Figure 1e** and **Suppl. Table 1**). For read alignment, demultiplexing, quality control, count matrices, and fragment file generation, we developed and made available TrESFlow, a dedicated NextFlow pipeline.

Upon profiling the Karpas-422 cell line with TrES-seq, we obtained 6,795 cells for H3K27ac, 7,730 cells for H3K27me3, and 5,804 for RNA, of which 5,252 cells exhibited high-quality signals for all three modalities and minimal cross-contamination between histone modalities (**Figure 1f**). Beyond cross-contamination, we compared Karpas-422 profiles obtained with TrES-seq with those obtained with uCoTargetX, scMTR-seq, and nano-CT, as well as with publicly available data from scMTR-seq, nano-CT, NTT-seq, and 10X Genomics multiome. In Karpas-422, TrES-seq generated H3K27ac and H3K27me3 profiles with a comparable or higher number of unique fragments compared to other methods (**Figure 1g**), and a higher fraction of reads in peaks (FRiP) (**Figure 1h**), indicating increased signal specificity. Similarly, TrES-seq RNA data exhibited higher numbers of both UMIs and detected genes per cell compared to the other methods (**Figure 1i**). Compared with publicly available datasets, TrES-seq data showed HPTM profiles of quality comparable to nano-CT and NTT-seq data, but superior to scMTR-seq (**Suppl. Figure 1f**), and gene expression data of superior quality to both scMTR-seq and 10X multiome (**Suppl. Figure 1g**).

Single-cell profiling methods for individual or dual HPTMs using single tagmentation have been largely optimized for H3K27ac and H3K27me3, while they typically cannot reliably be used to assay histone modifications that are enriched within large genomic regions, such as H3K9me3, because of the large fragments that are generated (>1 Kb) and Tn5 bias towards accessible chromatin^24^. To overcome this challenge, we modified the TrES-seq protocol to introduce a second tagmentation step, which allowed us to generate smaller fragments after the first tagmentation and, in this way, capture histone modifications that are enriched in large genomic regions (**Suppl. Figure 2a**). The TrESFlow pipeline was adapted to support both single and dual tagmentation experiments. We tested this modified version of TrES-seq to profile H3K9me3, H3K27me3, and RNA in Karpas-422 cells. The average FRiP score and number of unique fragments per cell were comparable to those obtained by single tagmentation (**Figure 1j**). H3K27me3 profiles generated with dual or single tagmentation were highly correlated, as well as with those generated by nano-CT, on the same cell line (**Figure 1k**). Similarly, pseudo-bulk H3K9me3 TrES-seq profiles were concordant with bulk ChIP-seq profiles obtained from ENCODE and were anti-correlated with both H3K27ac and H3K27me3 profiles from our independent experiments (**Figure 1k**). The RNA data obtained with TrES-seq were comparable with data obtained from the 10X Genomics scRNA-seq or bulk RNA data on the Karpas-422 cells (**Figure 1k**). Visual examination of germinal center B-cell lymphoma markers *MYC* and *BCL6* confirmed the enrichment of H3K27ac at their promoter concomitantly with high RNA abundance (**Figure 1l** and **Suppl. Figure 2b**). Conversely, broad H3K27me3 peaks were observed across the gene bodies of *IRF4* and *PRDM1*, which are markers of memory and plasma B cells, respectively, and are silenced in Karpas-422 (**Figure 1l** and **Suppl. Figure 2b**). Similarly, representative regions confirmed the specific enrichment of H3K9me3 in large genomic regions in a mutually exclusive manner with H3K27me3 and expressed genes (**Figure 1m** and **Suppl. Figure 2c**).

Lastly, the introduction of a sample barcode in the TrES-seq protocol enables multiplexing different samples in the same experiment. We tested this possibility by concurrently analyzing 5 different cell lines, including one immortalized human lymphoblastoid cell line (GM12878), two cell lines derived from Diffuse Large B Cell Lymphoma (DLBCL) patients (Karpas-422 and WSU-DLCL2), one cell line derived from a plasma cell leukemia patient (JJN3), and one murine lymphoma cell line (A20) (**Suppl. Figure 3a** and **Suppl. Table 2**). Cell lines were individually tagmented and labeled with a sample barcode and then mixed for triple barcode ligation to uniquely label each cell. Clustering of human cells based on individual modalities invariably identified well-defined clusters corresponding to individual cell lines, reflecting distinct transcriptional and epigenetic profiles (**Suppl**. **Figure 3b**). For each cell line, HPTM marker peaks show mutually exclusive signals for H3K27ac and H3K27me3 (**Suppl. Figure 3c**), consistent with the lack of cross-contamination (**Suppl. Figure 3d-e**). Overall, TrES-seq is a robust protocol that allows to profile RNA and two histone modifications with high specificity in the same cell, while implementing split-and-pool barcoding and sample multiplexing to reduce the costs of data generation.

### Multimodal profiles of primary tissue reveal multiple transcriptional states for the same H3K27me3 and H3K27ac profile

To test the TrES-seq protocol on human tissue samples and match the epigenetic and transcriptional status of different cell populations, we analyzed a human tonsil tissue obtained from a patient who underwent tonsillectomy. For this sample, we deep-sequenced the DNA modalities, reaching a saturation point for the number of unique fragment pairs for both H3K27ac and H3K27me3 (**Suppl. Figure 4a**). After demultiplexing and QC filtering (**Suppl. Figure 4b**), we obtained 9548 cells with high-quality RNA profiles (300-10,000 UMIs), among which 3053 cells also exhibited high-quality H3K27ac (300-15,000 unique fragments per cell) and 4181 cells high-quality H3K27me3 (300-20,000 unique fragments per cell) (**Figure 2a** and **Suppl. Figure 4c**). Cell typing was performed based on cell transcriptional profiles via label transfer^25^, using as a reference a previously published single-cell atlas including human tonsils^26^. Overall, we identified 15 distinct cell types, subtypes, and transitional cell states in this tissue (**Figure 2a** and **Suppl. Figure 4d**). Unbiased clustering of single-cell RNA profiles identified 14 stable clusters, which largely matched cell populations defined by label transfer (**Figure 2b**). In contrast, unbiased clusters based on either H3K27ac or H3K27me3 profiles recapitulated the cell types in our dataset (e.g. B vs T cells), but often did not discriminate among cell subtypes, such as naïve and memory B cells, or CD8+ and CD4+ T cells (**Figure 2b**). To robustly quantify these differences, we used different metrics to compare unbiased clusters with ground truth labels, such as Adjusted Rand Index (ARI) and conditional entropy (H), and we varied clustering parameters within a broad range of values. All metrics invariably confirmed that RNA-based clusters recapitulated cell types and subtypes with greater accuracy than histone modifications (**Figure 2c** and **Suppl Figure 4e**), which typically could not discriminate among distinct cell subtypes, irrespectively of the clustering resolution (**Suppl. Figure 4f**). These results were further confirmed in a subset of 1,880 cells with higher coverage (>600 RNA UMIs and >600 unique fragments in HPTM modalities) (**Suppl. Figure 5a-c**).

**Figure 2:**
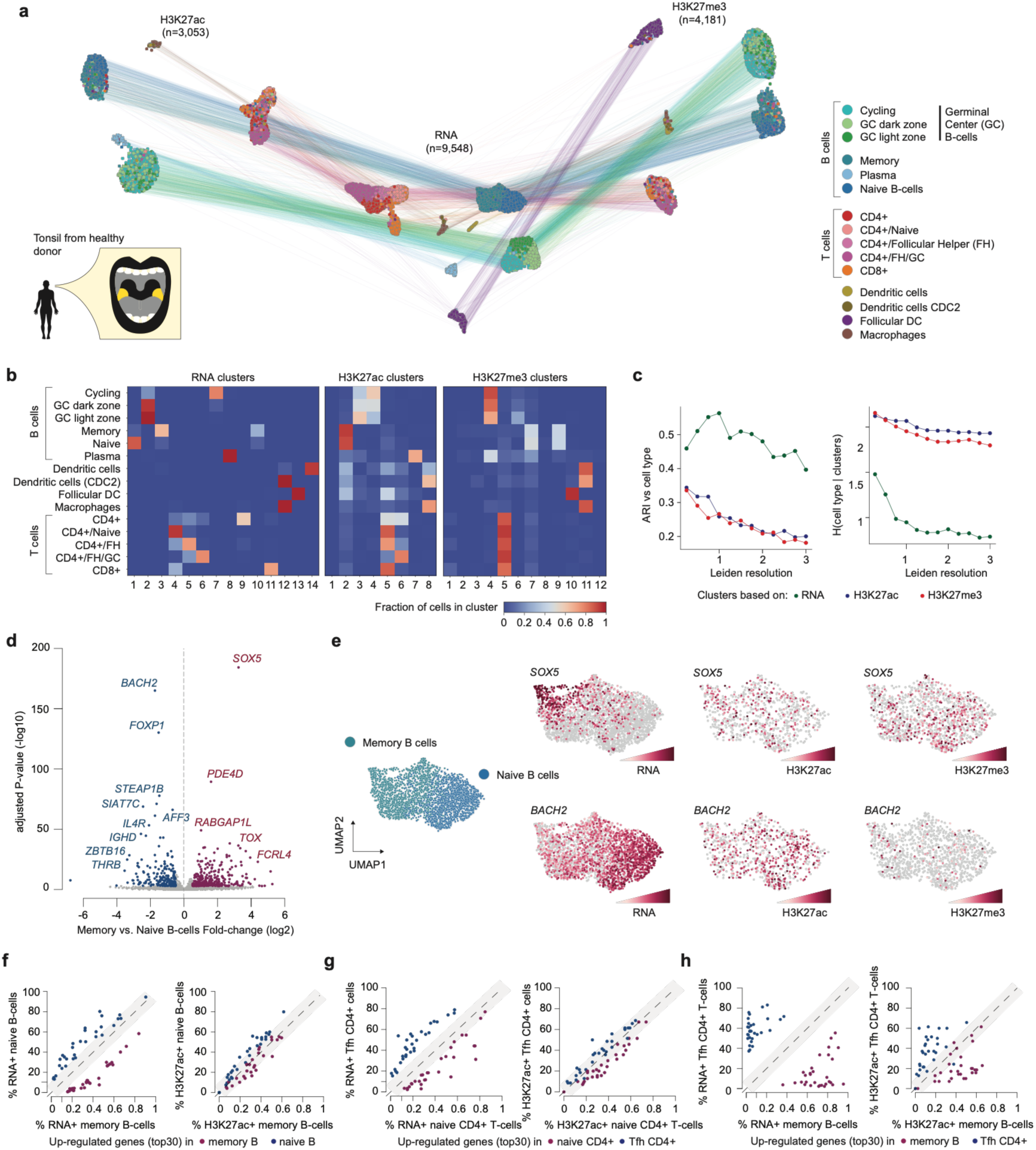
Joint epigenetic and transcriptional profiles in a primary tonsil tissue. **a.** Schematic representation of human tonsil and UMAP embeddings of the multimodal TrES-seq data for RNA (center), H3K27ac (left), and H3K27me3 (right). Corresponding cells in different modality UMAPs are connected by lines. Cells are color coded by the corresponding cell type. **b.** Confusion matrices (row normalized) comparing annotated cell types (rows) to unsupervised clusters (columns) for RNA (left), H3K27ac (middle) and H3K27me3 (right) at clustering resolutions. **c.** Adjusted Rand Index (ARI) between unbiased clusters and cell types (left) and conditional entropy of cell type given the clusters (right). **d.** Volcano plot showing differentially expressed genes between memory and naïve B cells. Selected differentially expressed genes are labelled. **e.** UMAP of naïve and memory B cells colored by cell type (left). The same embedding is colored by RNA expression, H3K27ac and H3K27me3 signal for SOX5 (top) and BACH2 (bottom). All signals are shown as log1p(CP10K). **f-h.** Scatter plots comparing the fraction of cells positive for RNA (left) or H3K27ac signal (right) between naïve and memory B cells (f), naïve and T follicular helper (Tfh) CD4⁺ T cells (g), and memory B cells and Tfh CD4⁺ T cells (h) for the top 30 upregulated genes in each cell type.

To assess epigenetic and transcriptional heterogeneity at specific genomic features, we focused first on cell subtype markers highlighted by differential expression analyses. For example, we identified several significantly differentially expressed genes between naïve and memory B cells (**Figure 2d** and **Suppl. Table 3**), including *SOX5*, the top upregulated gene in memory B cells, and *BACH2*, the top upregulated gene in naïve B cells (**Figure 2e**-left). Despite strong transcriptional differences, single cell H3K27ac and H3K27me3 signal intensities for these two genes were similar between the cell subtypes (**Figure 2e** – center, right). Similar results were obtained for the top differentially expressed genes between CD4+ naïve and CD4+ T-follicular helper cells (Tfh) (**Suppl. Figure 5d-e** and **Suppl. Table 3**). Notably, when comparing the percentage of positive cells in naïve and memory B cells (**Figure 2f**) or CD4+ naïve and Tfh cells (**Figure 2g**) for the top 30 differentially expressed genes, the numbers of RNA positive cells were clearly distinct among markers of the two cell subtypes (>10% difference), but more moderate (<10%) or no differences were found in the numbers of H3K27ac+ cells. Conversely, the same comparison between distinct cell types, such as memory B cells and CD4+ Tfh cells, showed high differences in the percentages of both RNA+ and H3K27ac+ cells between the two cell types (**Figure 2h**). Overall, matching gene expression, H3K27ac, and H3K27me3 data showed that, independently of the data resolution and number of clusters, cell types have distinguishable transcriptional and HPTM profiles, whereas transcriptionally different cell subtypes may exhibit similar H3K27ac and H3K27me3 profiles.

### H3K27ac at enhancers but not at promoters distinguishes between GC B cells and naive or memory B cells

To validate and further explore these results on a larger number of cells and samples, we generated TrES-seq data on tonsils obtained from three additional donors using either single-or dual-TAG TrES-seq protocols. We merged high-quality data from four different donors and obtained 27243 cells with RNA profiles, among which 11639 also exhibited high-quality data for at least one histone mark: H3K27ac (n= 4681 cells), H3K27me3 (n=10432 cells), and H3K9me3 (n= 3142 cells) (**Figure 3a** and **Suppl. Figure 6a-c**). Within this larger cohort, we confirmed that unbiased clustering of cells based on transcriptional profiles better recapitulated cell types and subtypes than H3K27ac, H3K27me3, or H3K9me3 profiles (**Suppl. Figure 6d-e**).

**Figure 3:**
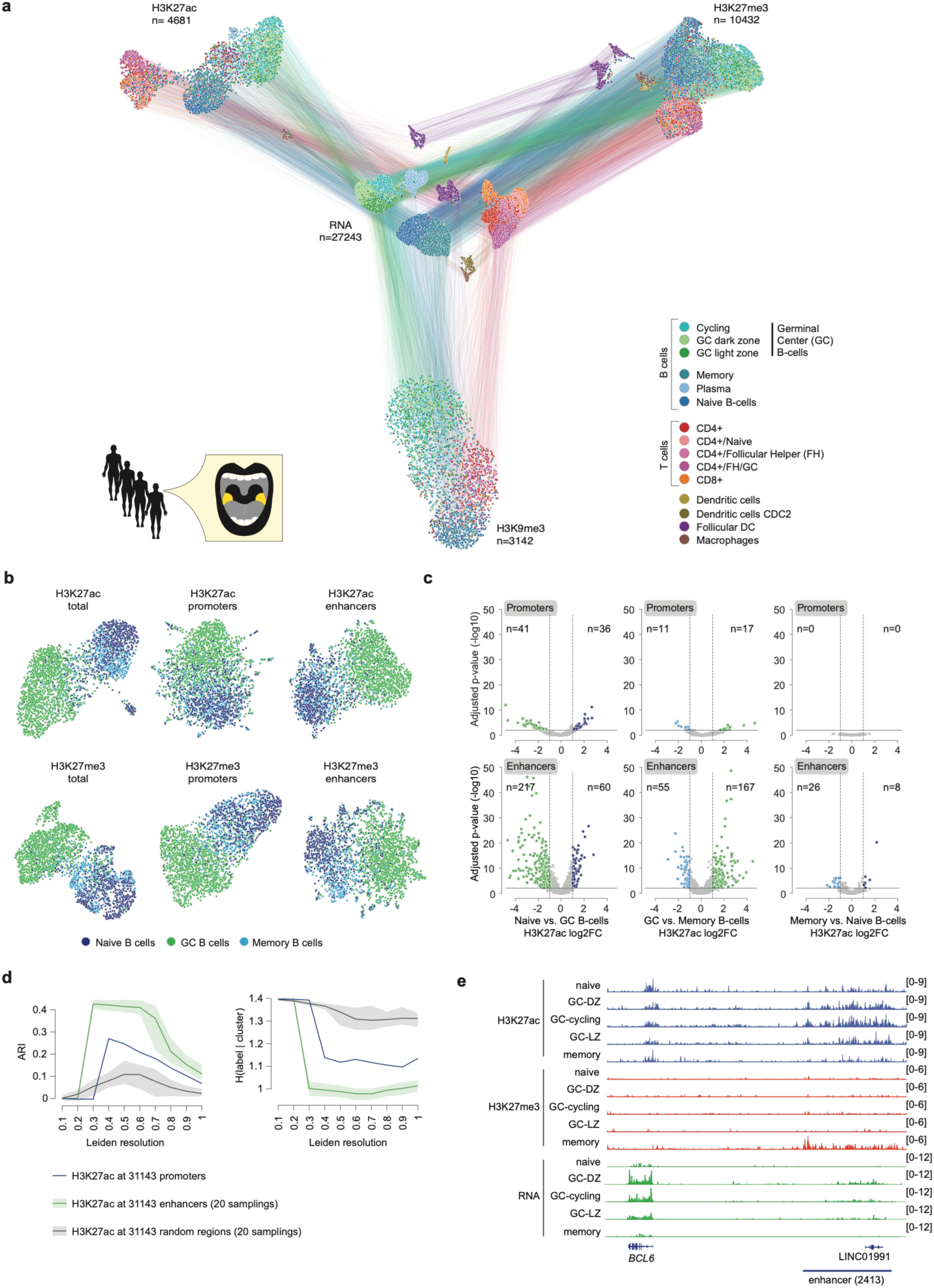
Epigenetic heterogeneity at enhancer and promoters in four primary tonsil tissuesa. UMAP of the multimodal TrES-seq data for RNA (center), H3K27ac (left), H3K27me3 (right), and H3K9me3 (bottom). Corresponding cells in different modality UMAPs are connected by lines. Cells are color-coded by the corresponding cell type. **b.** UMAP of naïve, germinal center (GC), and memory B cells based on H3K27ac (top) and H3K27me3 (bottom) in all tiles, promoter-overlapping tiles (TSS −1 kb, TSS +250 bp), and enhancer-overlapping tiles. **c.** Volcano plots of differential H3K27ac at promoters (top) and enhancers (bottom) across B-cell subtype pairs (naïve vs GC, GC vs memory, memory vs naïve). Points with −log10(adjusted P-value) > 2 and |log2 fold change| > 1 are colored, with counts (n) shown on each side. **d.** Comparison of unbiased clusters obtained with different resolution paramenters (X-axis) and cell type labels based on Adjusted Rand Index (ARI-left) and conditional entropy (H-right). Clusters were computed using size-matched sets of 31,143 tiles: all promoter-overlapping tiles (TSS −1 kb / +250 bp), random subsets of enhancer-overlapping tiles, and random genomic tiles. **e.** Genome browser views of matched TrES-seq profiles of aggregated signal tracks forH3K27ac, H3K27me3, and RNA at the *BCL6* locus and distal enhancer region for the indicated cell types.

Leveraging the higher number of cells in this dataset, we could distinguish H3K27ac and H3K27me3 signals at promoters and enhancers in B cell subpopulations^27^. Consistent with unbiased clusters based on total H3K27ac and H3K27me3 signals, H3K27ac at both promoters and enhancer regions was sufficient to distinguish germinal center (GC) B cells and naïve or memory B cell populations, although these populations seemed more distinct in the UMAP low-dimensional space when H3K27ac at enhancers was considered (**Figure 3b**). Differential H3K27ac analyses revealed more numerous and significantly different enhancers than promoters in naïve versus GC B cells and GC versus memory B cells, while no or only marginally significant differences were detected between promoters or enhancers of naïve and memory B cells (**Figure 3c** and **Suppl. Table 4**). These differences were further quantified by ARI and conditional entropy on a matching number of features and varying clustering resolution (**Figure 3d**).

Consistent with these results, a detailed analysis of *BCL6* promoter and enhancer regions in different B cell subpopulations confirmed a specific H3K27ac signal at the *BCL6* promoter across all B cell subtypes, even though *BCL6* mRNA expression was detectable only in GC B cells (**Figure 3e**). Gene expression differences were instead concordant with H3K27ac levels at enhancer regions, which exhibited higher H3K27ac in GC B cells than in naïve or memory B cells, with memory B cells exhibiting H3K27me3 in this region (**Figure 3e**). To verify that these results were independent of our single-cell technology, we analyzed publicly available H3K27ac ChIP-seq data of GC and naïve B cell subpopulations isolated from tonsils in the Blueprint project (https://ihec-epigenomes.org/research/projects/blueprint). In this independent data set, we confirmed an H3K27ac peak at the *BCL6* promoter in GC and naïve B cell subpopulations, but H3K27ac at the enhancer downstream of *BCL6* was observed exclusively in GC B cells (**Suppl. Figure 6f**). Thus, H3K27ac at enhancers but not at promoters matched transcriptional differences in GC versus memory and naïve B cells.

### Change of cell states is defined by modulation of gene expression within acetylated genes

So far, we showed a different degree of transcriptional and epigenetic heterogeneity between cell types and subtypes, with the latter often sharing similar epigenetic features, especially at gene promoters. To zoom in even further into distinct cell populations, we explored H3K27ac and H3K27me3 status in different cell states, among which a cell can dynamically and reversibly transition. For this purpose, we focused on cell state transitions among GC B cells. The GC is a specialized microenvironment of secondary lymphoid organs where naïve B cells undergo affinity maturation through multiple cycles of clonal selection, which results in the generation of memory B cells and plasma cells^28^ (**Figure 4a**). Specifically, within the GC, B cells proliferate and cycle between a “dark zone” (DZ) and a “light zone” (LZ)^29^, which are characterized by different transcriptional programs and cell interactions, consistent with distinct B-cell states^28^. Differential expression analysis between DZ and LZ cells in our tonsil dataset identified 1144 differentially expressed genes (|log_2_FC| > 0.5, FDR < 0.05) (**Figure 4b** and **Suppl. Table 5**). Conversely, differential H3K27ac analysis showed very few significant differences between these two populations, both at enhancer and gene promoters (**Figure 4c**), a result that was further confirmed for H3K27me3 and H3K9me3 (**Figure 4d**). Even when restricting the comparison to top differentially expressed genes, large RNA fold-changes corresponded to non-significant H3K27ac fold-changes (**Figure 4e**), indicating that the transition among these different cell states does not require altering H3K27ac and H3K27me3 in the cell. Strikingly, genes that were up-regulated in DZ compared to LZ cells were significantly enriched among genes that already exhibited high H3K27ac (top quartile) in LZ cells, and, similarly, genes up-regulated in LZ cells were enriched among the top quartile of H3K27ac+ genes in DZ cells (**Figure 4f**). This result suggested that cell state transition may often involve transcriptional modulation within a pool of H3K27ac+ genes.

**Figure 4:**
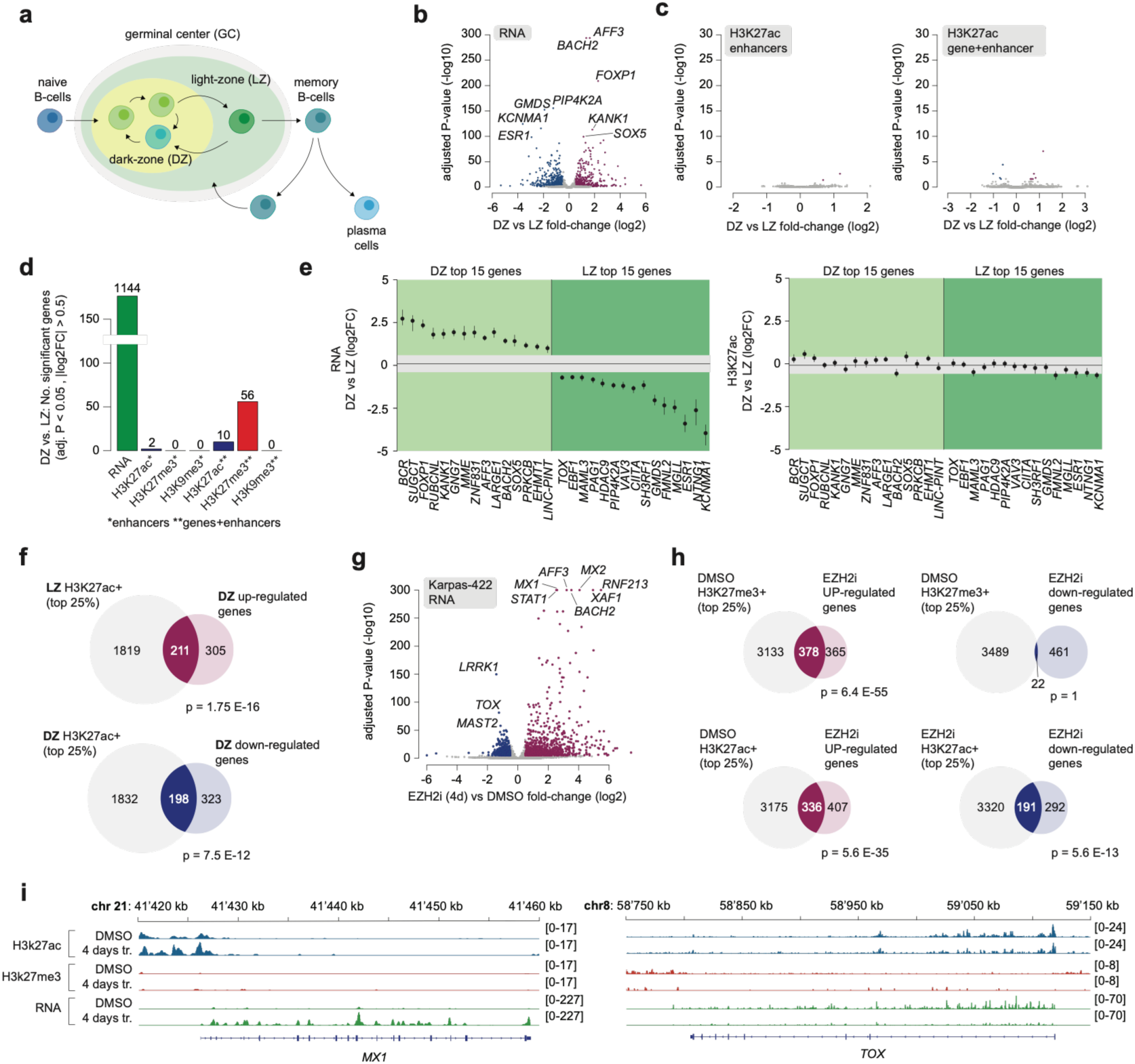
Joint epigenetic and transcriptional profiles of transitional cell states. **a.** Schematic representation of B-cell state transitions during the germinal center (GC) reaction.**b,c.** Volcano plots comparing RNA expression (b) and H3K27ac fragment abundance (c) between DZ and LZ B cells. In c, fragments were counted separately for each enhancer (left) or summed across each gene body and its linked enhancers (right). **d.** Numbers of genes exhibiting differential RNA expression or histone marks between DZ and LZ (adjusted P < 0.05, |log₂ fold change| > 0.5). Single and double asterisks denote enhancers alone and gene bodies plus enhancers, respectively. **e.** RNA (left) and H3K27ac (right) fold changes (log2) for the top 15 differentially expressed genes in each cell state (DZ and LZ). Points show median DZ/LZ log2 fold changes; bars span the 10th–90th percentiles of paired subset estimated across 20 random partitions. Gray bands mark ±0.5 log2 fold change. **f.** Overlap of genes in the top H3K27ac quartile in LZ with genes upregulated in DZ (top), and in top DZ quartile with genes downregulated in DZ (bottom). P-values are computed by one-sided Fisher’s test. **g.** Volcano plot showing differentially expressed genes between Karpas-422 cells treated with EZH2 inhibitor and DMSO. **h.** Overlap of genes upregulated (left) or downregulated (right) after EZH2 inhibition with genes in the top quartile of H3K27me3 in DMSO (top), H3K27ac in DMSO (bottom left) or H3K27ac after EZH2 inhibition (bottom right). **i.** Genome browser views of matched TrES-seq profiles of aggregated signal tracks for H3K27ac, H3K27me3, and RNA at the *MX1* and *TOX* loci for the indicated conditions.

To validate this result in an independent system, we compared epigenetic and transcriptional profiles of Karpas-422 cells before and after treatment with a selective EZH2 inhibitor (EZH2i). Briefly, Karpas-422 cells harbor an oncogenic mutation in the *EZH2* gene, which encodes for the catalytic subunit of the polycomb repressor complex 2 (PRC2). This mutation results in hyperactivation of EZH2 and increased H3K27me3.

Treatment with EZH2i ablates H3K27me3 and pushes the cell into a more differentiated but reversible cell state^30^. Hence, this model provided the opportunity of using TrES-seq to assess the association between transcriptional and epigenetic changes in cell state transitions induced by epigenetic perturbation. Karpas-422 cells were treated for 96 hours and profiled with TrES-seq, obtaining 6003 cells (4,974 for control and 1029 for treated conditions) with high-quality RNA, H3K27ac, and H3K27me3 profiles. As expected, H3K27me3 was significantly reduced genome-wide in treated vs. control cells (**Suppl. Figure 7a-c**). Differential expression analysis between treated and control cells revealed multiple significant transcriptional changes (**Suppl. Table 5**), especially among up-regulated genes, consistent with the depletion of the repressive histone mark H3K27me3. Indeed, H3K27me3+ genes in control cells were significantly enriched in those that were transcriptionally up-regulated upon treatment and depleted among those that were down-regulated (**Figure 4h**). Strikingly, beyond transcriptional changes induced by H3K27me3 depletion, we observed a significant enrichment of up-regulated and down-regulated genes among H3K27ac+ genes before and after treatment, respectively (**Figure 4h**). Indeed, a detailed analysis of the genomic region of one of the top upregulated (*MX1*) and downregulated genes (*TOX*) in EZH2-treated cells confirmed the presence of H3K27ac peaks at promoters in both control and treated cells (**Figure 4i**). This result demonstrated that even in the presence of a selective epigenetic perturbation, cell state transition modulated transcriptional activity of genes that exhibited and retained an active epigenetic state.

Overall, matching multimodal profiles of HPTM and gene expression in single cells suggest a model where distinct cell types exhibit different transcriptional and epigenetic profiles; transcriptionally different cell subtypes may share instead similar epigenetic features, with a different degree of heterogeneity at promoter and enhancer regions, and, lastly, transitions among cell states largely rely on transcriptional modulation of genes that are already in an epigenetically active state.

## Discussion

Single-cell profiling of HPTMs and gene expression has the potential to define how specific histone marks relate to transcriptional heterogeneity in complex tissues, and how their interplay defines cell identity. However, only a limited number of studies have used such multimodal sequencing approaches. The implementation of methods such as nano-CT, uCoTargetX, or scMTR-seq requires extensive optimization, even when protocols and commercial kits are available. Moreover, to increase the number of HPTMs that are assayed in each experiment, existing technologies often rely on pre-assembled antibody–pA-Tn5 complexes, at the expense of HPTM signal specificity and accurate interpretation of the results.

By developing TrES-seq, we sought to design a modular technology that allows to generate high-quality data while being easy to test and implement in a cost-effective manner. Importantly, while TrES-seq showed superior performance compared to competing technologies, the flexibility of this protocol will allow further improvement in its throughput and applicability to map other HPTMs, chromatin architectural proteins, and transcription factor bindings. However, the availability of species-specific nanobodies, which are essential to limit cross-contamination, defines the number of multimodal proteins that we can simultaneously detect. Currently, mouse-and rabbit-specific nanobodies enable the concurrent detection of two HPTMs with high specificity, but the development of additional species-specific nanobody-Tn5s is required to increase the number of HPTMs that can be assayed without compromising data quality. Furthermore, bivalent regions that are simultaneously marked by two HPTMs cannot be simultaneously detected using Tn5-based methods, since the two antibody-Tn5-nanobody complexes will compete and cut the same region. To facilitate the adoption of the TrES-seq protocol, we designed a multiplexed barcoding strategy, which is independent of microfluidic technologies and, after optimization, it reduced the cost for obtaining high-quality single-cell RNA and HPTM data to ∼1000-1500 USD per experiment. Lastly, data generation is only useful if accompanied by reliable and streamlined data analyses. To this purpose, we developed TrESFlow, a Nextflow pipeline that automates data processing and quality control across modalities. By integrating these steps into a single efficient workflow, TrESFlow reduces the need for custom scripting, enabling robust, standardized, and reproducible procedures across experiments.

To demonstrate the potential of TrES-seq, we examined a hierarchy of cell identities in human tonsils, as a representative example of complex tissue. Here, we determined different degrees of heterogeneity for RNA and histone mark profiles across cell types, subtypes, and cell states, as well as between distinct genomic regions (i.e. promoters and enhancers). Whereas these associations will need to be further tested in other tissues and coupled with functional studies to assess their generalizability, in tonsils we showed that transitions among cell states do not require epigenetic reprogramming, while different cell types exhibit distinctive epigenetic features. Recently, several single-cell atlases based on scRNA-seq have identified multiple known and new cell subpopulations. In this context, joint epigenetic and transcriptional profiles will help discriminate between transitioning cell states and cell types or subtypes.

Moreover, our analyses of either naturally occurring or treatment-induced cell state transitions revealed broad transcriptional changes occurring among genes that already exhibited and retained an epigenetically active state, suggesting epigenetic priming and memory. Similar results were reported for stimulated macrophages and T cells where several induced genes already exhibited active histone marks and promoter-bound RNA polymerase II before induction^31^, or in hematopoiesis, where formation of de novo enhancers was shown to anticipate transcriptional activation and prime lineage commitment^32^. However, most studies have so far relied on bulk measurements of epigenetic and transcriptional profiles, with a few notable exceptions where chromatin accessibility was again shown to precede transcriptional activation^15^. In this context, single-cell multimodal datasets generated from longitudinal experiments are needed to assess response dynamics to cell perturbation, pinpoint epigenetic features that can predict transcriptional response, and estimate the extent of molecular changes that are required to reprogram cell identity. This will be particularly relevant in malignant conditions, such as cancer, where loss of lineage fidelity and trans-differentiation are common mechanisms of therapeutic resistance. Overall, new multimodal technologies, such as TrES-seq, enable high-throughput single-cell multimodal data generation across multiple contexts, and, when coupled with functional experiments, will be key to determine how heterogeneous chromatin states influence transcription and, thus, cell identity and function.

## Author contribution

Experimental Data Generation: V.K.P., N.K, A.S.M., and J.E; Data Analyses, A.A, D.S. and G.C.; Sample collection K.L; Writing, E.O. and G.C., with contributions from all authors; Conceptualization, E.O. and G.C.; Supervision, E.O. and G.C.

## MATERIALS AND METHODS

### Cell lines and cell culture

DLBCL cell lines WSU-DLCL2 and Karpas-422 were obtained from DSMZ, lymphoblastoid cell line GM12878 and multiple myeloma JJN3 from a previous collaboration, and mouse DLBCL cell line A20 was obtained from ATCC. Karpas-422, WSU-DLCL2, GM12878 were maintained in suspension in 10 cm tissue culture dishes (TPP, 93100) in RPMI-1640 medium (Thermo Fisher Scientific, 11875093) supplemented with 10% FBS (Thermo Fisher Scientific, 10270106) and 50 U ml−1 penicillin-50 µg ml−1 streptomycin (Thermo Fisher Scientific, 15070063). JJN3 was maintained in culture vertically in T75 flasks (Bio-One, 658175) in IMDM (Thermo Fisher Scientific, 31980048) and DMEM (Thermo Fisher Scientific, 41966029) supplemented with 10% fetal bovine serum (FBS) (Thermo Fisher Scientific, A5256701), and 50 U ml−1 penicillin-50 µg ml−1 streptomycin (Thermo Fisher Scientific, 15070063).A20 cells were maintained in suspension in 10 cm tissue culture dishes (TPP, 93100) in RPMI-1640 + GlutaMAXTM (Thermo Fisher Scientific, 61870010) supplemented with 10% heat-inactivated fetal bovine serum (FBS, heated 30 min at 56°C) (Thermo Fisher Scientific, A5256701), 1 mM sodium pyruvate (Sigma-Aldrich, S8636), 10 mM HEPES (Sigma-Aldrich, H0887), 5 mM d-glucose (Sigma-Aldrich, G8769), 1% MEM non-essential amino-acids (100x) (Thermo Fisher Scientific, 11140050), 25 µg/mL gentamicin (Sigma-Aldrich, G1397) and 50 μM 2-mercaptoethanol (Thermo Fisher Scientific, 31350010). Cells were maintained in a sterile, humidified incubator at 37°C and 5% CO_2_ and sub-cultured every 3 days to maintain adequate cell confluency.

### Patient sample collection and pre-processing

Adult patients scheduled for tonsillectomy at CHUV for either sleep apnea or recurrent tonsillitis were informed about the study and invited to participate. After obtaining informed consent, fresh tonsil samples were collected in 50 ml Falcon tubes containing 30 mL of cold RPMI supplemented with 100 μg/ml primocin and transported to the laboratory. The tissue was mechanically dissociated, passed through a 70 μm strainer and washed twice with cold PBS. Samples were then incubated with red blood cell lysis buffer (Santa Cruz Biotechnology) according to the manufacturer’s instructions, washed with PBS, and further processed as described below. All procedures were carried in accordance with the Ordinance on human research and were previously approved by the *Commission cantonale d’éthique de la recherche sur l’être humain* (CER-VD, project-ID2023-01787).

### TrES-seq

#### 1. Tn5 Assembly

For assembled nanobody-Tn5 for single tagmentation, equal volume of barcoded 100 µM Nano_AdapA_BcA, Nano_AdapA_BcB modality oligonucleotides and 100 µM Tn5_AdapB_Ligation oligonucleotide were dispensed into separate tubes. To each tube, equal volume of 100µM Me_Rev_pMENTs_Blocked ME was added, followed by thorough mixing. The oligonucleotide mixtures were annealed using the following thermocycler program: 95°C for 5 minutes, followed by a gradual ramp-down to 20°C at 0.1°C/s. Then the following components were combined: annealed barcoded Nano_AdapA_BcA_X (mouse specific) or Nano_AdapA_BcB_X(rabbit-specific)/ Me_Rev_pMENTs_Blocked ME, annealed Tn5_AdapB_Ligation / Me_Rev_pMENTs_Blocked ME, 100% glycerol, 2x Dialysis Buffer, 20% glycerol, and Nanobody-Tn5 enzyme (targeting mouse or rabbit, respectively). The components were gently mixed and incubated at room temperature for 1 h to enable efficient enzyme loading and stored at −20°C.

#### 2. Ligation plate preparation

Combinatorial barcoding was performed using three sets of oligonucleotides (**Suppl. Table 1**) which were annealed as follows. For ligation-1, 2.62 µL of each 100µM ligation-1 barcodes oligos were distributed into 48 individual wells of a 96-well plate. To each well, 2.42 µL of 100µM linker-1 and 16.95 µL of nuclease-free water were added. For ligation-2, 2.62 µL of each 100µM ligation-2 barcodes were combined with 2.42 µL of linker-2 and 16.95 µL of nuclease-free water in each well of a 96-well plate. For ligation-3, 2.62 µL of each ligation-3 barcodes oligos were mixed with 2.42 µL of linker-3 and 16.95 µL of nuclease-free water. Then oligonucleotides were annealed by incubating at 95°C for 5 minutes, followed by a gradual ramp-down to 20°C at a 0.1°C/s using a thermocycler. Following annealing, 10µL from each well was transferred to two new 96-well plates. The plates were sealed and stored at –20°C until further use.

#### 3. Antibody & Nanobody Incubation

100,000 cells per sample were aliquoted in different DNA LoBind Tubes. Samples were centrifuged at 500x g for 5 minutes at 4°C, and the supernatant was aspirated. To each cell pellet, antibody buffer containing H3K27me3 mouse antibody and H3K27ac rabbit antibody were added. Subsequently, mouse-specific nanobody-Tn5 (single or dual tagmentation) and rabbit-specific nanobody-Tn5 (single or dual tagmentation) were added to each sample. Samples were mixed gently and incubated overnight at 4°C on the horizontal roller for antibody and nanobody Tn5 binding.

#### 4. Tagmentation

After overnight incubation with the antibodies and nanobody Tn5s, samples were centrifuged at 500x g for 5 minutes at 4°C, and the supernatant was aspirated. Cell pellets were resuspended in Dig Wash buffer and centrifuged again 500x g for 5 minutes at 4°C. Washing step was repeated two times. Cells were then resuspended in tagmentation buffer and incubated the mixture at 37°C for 1 hours to perform tagmentation.

#### 5. Reverse transcription

Following tagmentation, samples were centrifuged at 500 x g for 5 minutes at 4°C and washed two times with NIB buffer. The final pellet was resuspended in NIB buffer and combined with reverse transcription (RT) mix and RT primers (**Suppl. Table 1**). Reverse transcription was performed in a thermocycler. After reverse transcription, all 12 samples were pooled and added NIB buffer was added. The pooled sample was centrifuged at 500 x g for 5 minutes at 4°C, and the supernatant was removed. A final two washes were performed with NIB before combinatorial barcoding.

#### 6. *In situ* combinatorial barcoding

Ligation plates were thawed at room temperature before use. To the reversed transcribed samples, 1x NEBuffer3.1, 10x T4 DNA ligase buffer, 20 mg/mL BSA, 10x NEBuffer 3.1, Enzymatics RNase Inhibitor, T4 DNA ligase was added and samples were mixed thoroughly. The nuclei with ligation mix were added to each well of the ligation plate.

Plates were incubated at 37°C with gentle shaking at 350 rpm for 1 hour to enable ligation. Following ligation-1, blocking oligo-1 was added to each well, and the plate was incubated again at 37°C at 350 rpm for 30 minutes. Then, all wells were pooled, and the sample was centrifuged at 1000 x g for 10 minutes at 4°C. The supernatant was carefully aspirated.

The second and third ligation rounds of combinatorial barcoding were performed using the same procedure as the ligation-1, each followed by incubation with blocking oligo-2 respectively. After the final blocking step (blocking oligo-3), the sample was washed twice with NIB buffer. Nuclei were then counted. 10,000 nuclei were transferred into Eppendorf tubes. To each tube, 2x Reverse crosslinking buffer, Proteinase K and SUPERase-In were added. Samples were mixed thoroughly and incubated at 55°C for 2 hours with shaking at 800 rpm. Following incubation, phenylmethylsulfonyl fluoride (PMSF) was added to each tube to inactivate Proteinase K. Samples were then incubated at room temperature for 10 minutes. Processed sub libraries were either used immediately for next steps or stored at –80°C for future steps.

#### 7. DNA and RNA separation

Dyna MyOne C1 beads were activated as follows: beads were transferred to a clean tube and washed three times with 1x B&W-T buffer. A final wash was performed using 1x B&W-T/RI. Beads were then resuspended in 2x B&W/RI buffer. In parallel, sub-libraries were thawed on ice. Activated bead suspension was added to the thawed sub-library and mixed thoroughly. The mixture was incubated at room temperature for 1 hour on a rotating platform to enable RNA/cDNA hybrids to bind with the Dyna MyOne C1 beads. After incubation, tubes were placed on a magnetic rack to separate the bead-bound fraction from the supernatant. The supernatant, containing DNA, was carefully transferred to a new tube. The magnetic bead pellet was retained, containing the captured RNA/cDNA hybrid complexes.

#### 8. DNA Library Preparation for single tagmentation

Supernatant containing DNA in fresh tube was purified using the Zymo DNA Clean & Concentrator kit following manufacturer’s protocol. DNA was eluted in nuclease-free water. The purified DNA was then amplified. Amplification was performed amplification in a thermocycler. Amplified libraries were subjected to custom paired-end sequencing with a target depth of approximately 100 million reads per library.

#### 9. RNA library preparation

Bead-bound RNA/cDNA hybrids were washed three times with 100 µL of 1x B&W-T/RI buffer, followed by a single wash with STE/RI buffer. The beads were then resuspended in 50µL of freshly prepared Template Switch and incubated for 30 minutes at room temperature on a rotating platform to enable efficient template switching. This was followed by a 90-minute incubation at 42°C with shaking at 350 rpm, during which the reaction was briefly mixed by pipetting every 30 minutes. After incubation, 100µL of nuclease-free water was added to the tube, and beads were magnetically separated to discard the supernatant. Then beads were washed once with 1x STE buffer, and the supernatant was aspirated. The bead pellet was resuspended in cDNA PCR mix and transferred to a clean PCR tube for pre-amplification. Pre-amplification was carried out and amplified cDNA was purified using 0.8xSPRI beads cleanup and eluted in nuclease-free water. The cDNA was quantified by Qubit dsDNA quantification kit. 50ng of purified pre-amplified cDNA was fragmented by incubating with tagmentation mix. Then the mixture was incubated for 30 minutes at 37°C. The mixture was then purified with Zymo DNA clean and concentrate kit and eluted. The tagmented fragments were amplified in a thermocycler. Amplified libraries were subjected to custom paired-end sequencing to a target depth of approximately 100 million reads per library.

### scRNA-Seq

scRNA-seq was performed using a Chromium Next GEM Single Cell 3′ kit (10x Genomics) following the manufacturer’s protocol. Libraries were sequenced on a HiSeq 4000 system (Illumina).

### Nano-CUT&Tag library preparation

Nano-CUT&Tag (nano-CT) was performed in Karpas-422 cells following the protocol described in^18^. Briefly, cells were harvested, washed in ice-cold PBS, and permeabilized. Permeabilized cells were incubated with different species primary antibodies and species-specific nanobody-Tn5s. Antibody and nanobody-Tn5 binding reactions were carried out on a horizontal roller at 4 °C overnight. Subsequently, cells were tagmented at 37 °C for 1 hour. After tagmentation, nuclei were processed using the Chromium Next GEM Single Cell platform (10x Genomics) according to the manufacturer’s instructions. Following GEM generation and barcoding, emulsions were subjected to post-GEM incubation and subsequent cleanup to recover barcoded DNA. Purified DNA was quantified using Qubit. Libraries were then subjected to tagmentation and final library amplification. Library quality and fragment size distribution were assessed prior to sequencing. Sequencing was performed on an Illumina platform using paired-end reads according to standard nano-CT specifications.

### EZH2 inhibitor treatment

Karpas-422 cells (1 × 10⁶) were seeded in RPMI medium supplemented with 10% fetal bovine serum (FBS) and 1% penicillin–streptomycin and treated with 0.5 µM EZH2 inhibitor EPZ-6438 (tazemetostat) or dimethyl sulfoxide (DMSO) as a vehicle control for 96 h. Following treatment, cells were collected and processed for TrES-seq. Sequencing reads were aligned to hg38 reference genome.

## DATA ANALYSIS

### Data preprocessing

Raw TrES-seq sequencing data were processed with TrESFlow, a Nextflow pipeline that integrates demultiplexing, barcode error correction, read trimming, alignment, and deduplication.

### Read trimming and FASTQ splitting

Tagged reads were trimmed with TrimGalore v0.6.10 (minimum base quality 10; minimum retained length 20 nt; paired-end synchronization). Trimmed reads were split into FASTQs by sample, condition, and mark using the sample and modality barcodes.

### Alignment and deduplication

DNA FASTQs were aligned to the appropriate reference genome (hg38/GRCh38 or GRCm39) using BWA-MEM2 v2.2.1 with exclusion of blacklisted regions^33^. PCR duplicates were marked with GATK MarkDuplicates v4.6.0.0 using CB as the barcode field and removed with samtools v1.21 based on the duplicate flag (0×x400). BWA-aligned, deduplicated, cell-tagged BAM files were used downstream. RNA FASTQs were aligned with STARsolo v2.7.11b in two-pass mode using junction-based filtering and removal of non-canonical introns; CB/UMI processing used cell barcodes from the SAM CB tag, UMI deduplication allowed one mismatch, and cell calling used the EmptyDrops-based filter. STARsolo gene-count matrices were used downstream.

### RNA modality

STARsolo gene-by-cell matrices were loaded into Scanpy and per-cell QC metrics were computed. Cells were retained using dataset-specific thresholds (**Suppl. Table 6**). RNA counts were library-size normalized, log1p-transformed, highly variable genes were selected (Seurat v3 flavor; 2,000 genes), PCA was computed, a kNN graph was built (cosine distance), and UMAP embeddings were generated. For analyses requiring paired measurements, cells were restricted to the exact barcode intersection across modalities (RNA ∩ H3K27ac ∩ H3K27me3), and each modality was subset to the shared cell set.

### Cross-mark signal profiles

For each dataset, fragments from cells passing quality control were pooled separately for H3K27ac and H3K27me3. Peaks were called with MACS3 as described above, using a q-value cutoff of 10⁻⁴. Peaks whose intervals plus 5-kb flanks intersected ENCODE blacklisted regions were excluded. Pseudobulk coverage was normalized separately for each mark per million retained fragments using MACS3--SPMR. Using deepTools v3.5.5, coverage for both marks was summarized in 100-bp bins within ±5 kb of the midpoints of the same selected peaks, using each dataset’s own peaks for the indicated mark as anchors.

### Signal comparison

Genome-wide signal tracks were generated from BAMs using deepTools v3.5.5 bamCoverage with RPGC normalization (bin size 20). Track similarity was quantified using multiBigwigSummary (bins mode; 10 kb bins) followed by Spearman correlation and visualization with plotCorrelation. Aggregated coverage tracks were visualized in IGVtools v2.19.7. Single-cell-level read/fragment stacks were generated from BAMs using Bioconductor tooling and plotted with ggplot2. For histone marks, paired-end fragments were reconstructed and plotted as segments spanning fragment start-end. For visualization, cells were subsampled per track to a target of 200 cells by selecting those with the highest number of locus-overlapping segments/fragments.

### Promoter-and peak-centered metagene plots

H3K27ac and H3K27me3 coverage was RPGC-normalized with deepTools v3.5.5 bam Coverage. For the top 50 marker genes per human cell line, compute Matrix quantified coverage in 100-bp bins within ±20 kb of transcription start sites, preserving input order. Coverage for both marks was quantified in 20-bp bins within ±5 kb of the midpoints of blacklist-filtered H3K27ac H3K27me3 peaks. Mean profiles and heatmaps were generated with plotProfile and plotHeatmap.

### Tonsil data processing

Raw TrES-seq reads from four tonsil donors were processed with TrESFlow. Quality filters were applied by data type (RNA or HPTMs) and low-quality clusters were removed based on a signal score based on unique fragments × fraction of fragments in peaks (FRiP). Doublet filtering used DoubletDetection v4.3.0 calls for RNA and SnapATAC2 calls for DNA. RNA labels were transferred from a scRNA-seq reference dataset of Tonsil tissue (GSE269695) using scvi-tools 1.5.1 and propagated directly to matching DNA profiles using cell barcodes. Analyses used Tonsil1, Tonsil1 Highly Covered (>600 RNA UMIs and unique DNA fragments per cell, independently by modality) and the merged four-donor dataset. UMAPs used nearest-neighbor graphs built from RNA principal components or DNA spectral embeddings tile counts (1kb for H3K27ac, 10kb for H3K27me3 and H3K9me3). The donor-merged DNA datasets used Harmony correction (harmonypy v2.1.0) for single-versus dual-tagmentation. RNA remained uncorrected.

### Cluster agreement

Unbiased Leiden clusters were identified at multiple resolutions based on the nearest-neighbor graphs built from RNA principal components or DNA spectral embeddings. For each tonsil dataset, tri-profiled cells were retained, and modality and Leiden resolution cluster assignments were cross-tabulated against RNA labels. Agreement was quantified using the adjusted Rand index (ARI).

### Differential gene expression/marking analysis

Raw RNA counts were summed by gene within three disjoint random cell subsets per population to generate pseudobulk replicates. Pairwise differential expression was tested with DESeq2 using cell population as the design factor. For visualization, genes with FDR ≤ 0.001, absolute log₂ fold change ≥ 0.5 and RNA detection in ≥1% of cells in the population with higher expression were ranked by FDR, retaining up to 30 per direction. Mega-enhancer regions (MERs) from^27^ were lifted to hg38 and linked to genes within 100 kb. SnapATAC2 counted H3K27ac fragment overlaps at individual MERs and over the union of each gene body (±1 kb) with its linked MERs. We randomly divided cells in each zone into three non-overlapping subsets of similar size and summed raw H3K27ac fragment counts within each subset. We compared the resulting pseudo bulks using DESeq2 with zone as the model term, poscounts size factors, and Wald tests. Differential H3K27ac required a Benjamini–Hochberg FDR ≤ 0.001 and an absolute log₂ fold change ≥ 0.5.Effect sizes were recalculated for paired subsets across 20 random partitions, with bars spanning the 10th–90th percentiles of the resulting estimates.

### Cross-modality visualization

Histone measurements were displayed on the RNA UMAP using exact cell matches. RNA counts and histone gene body counts by SnapATAC2 were scaled to 10,000 per cell. Cells were colored by log(1 + normalized signal), with color scales capped at the 99th percentile of positive values for each gene and modality.

### Comparison of percentages of positive cells

The percentage of positive cells for each modality was computed as the fraction of cells with non-zero count for that modality. For RNA, it corresponds to non-zero gene counts, for DNA modalities at gene level, it corresponds non-zero count in a window corresponding to the gene body ± 1kb (**Figure 2f-h**).

### UMAP visualization of promoter and enhancer epigenetic features

Analyses were restricted to non-plasma B cells with histone counts (2,635 cells for H3K27ac and the matched H3K27me3 library). Stages were defined from RNA labels as naïve, germinal-center (cycling, dark-zone, and light-zone), and memory. Chromatin features were 1 kb genome tiles for H3K27ac and 10 kb tiles for H3K27me3; tiles with no count in the analyzed cells were dropped. For each mark, three feature sets were compared: all tiles with a count; promoter tiles overlapping TSS − 1 kb to TSS + 250 bp; and enhancer tiles overlapping B-cell mega-enhancer regions^27^ (MERs, lifted to hg38). Each feature set was embedded with SnapATAC2 cosine spectral embedding, Harmony-corrected for tonsil, then used to build a 15-nearest-neighbor graph (Euclidean) and UMAP (min_dist = 0.3, spread = 1). UMAPs were colored by B cell stage.

### Differential H3K27ac at promoters and enhancers

Stage-specific H3K27ac was tested on the same 2,635 cells for three pairwise contrasts: naïve vs GC, GC vs memory, and memory vs naïve. Enhancers were MER intervals; each cell’s enhancer count was the sum of overlapping 1 kb H3K27ac tiles. Promoters were TSS − 1 kb to TSS + 250 bp, likewise summed over overlapping tiles. Features detected in fewer than 5% of cells in both groups of a contrast were excluded. Differential abundance was tested with snapatac2.tl.diff_test (earlier stage as group 1; min_pct = 0.05, min_log_fc = 0). Features with -log_10_(adjusted *P*) > 2 and |log₂ fold change| > 1 were called stage-specific; volcano plots show all tested features, with non-significant points in grey.

### Size-matched clustering of promoters, enhancers, and random tiles

To separate annotation effects from feature-set size, H3K27ac clustering was compared at a fixed tile count equal to the promoter set (31,143 tiles). Promoters used the full set once. Enhancer tiles and genome-wide random tiles (drawn from all tiles with a count) were subsampled without replacement to the same size across independent draws. Each draw was re-embedded, Harmony-corrected, and clustered with Leiden over resolutions 0.1–1.0 (step 0.1). Agreement with stage was scored as adjusted Rand index (ARI) and conditional entropy *H*(stage | cluster). Curves report the mean across draws, with the shaded band the range of those draws.

## Data availability

The following publicly available datasets were used in this study: GSE269695 -scRNA-seq reference dataset of Tonsil tissue; 10x Genomics Multiome Human PBMCs (https://www.10xgenomics.com/datasets/10-k-human-pbm-cs-multiome-v-1-0-chromium-x-1-standard-2-0-0); GSE277856 -Multimodal sequencing of hPSCs (scMTR-seq); GSE220193 -Multimodal sequencing of K562 cells (uCoTargetX: Bulk Coverage tracks); GSE212588 -Multifactorial chromatin profiling of K562 cells (NTT-seq); GSE198467 -Multimodal profiling of two histone modifications of Mouse Brain (nano-CT); and from the ENCODE portal (https://www.encodeproject.org/), Bulk-RNA and H3K9me3 ChIP-seq of Karpas-422 cells, with identifiers ENCSR314LXG and ENCSR978BQI, respectively.

