## Supplementary Data for "TrES-seq integrates single-cell RNA and nanobody-based histone mark profiles to decode epigenetic and transcriptional axes of cell identity"

Suppl. Figure 1

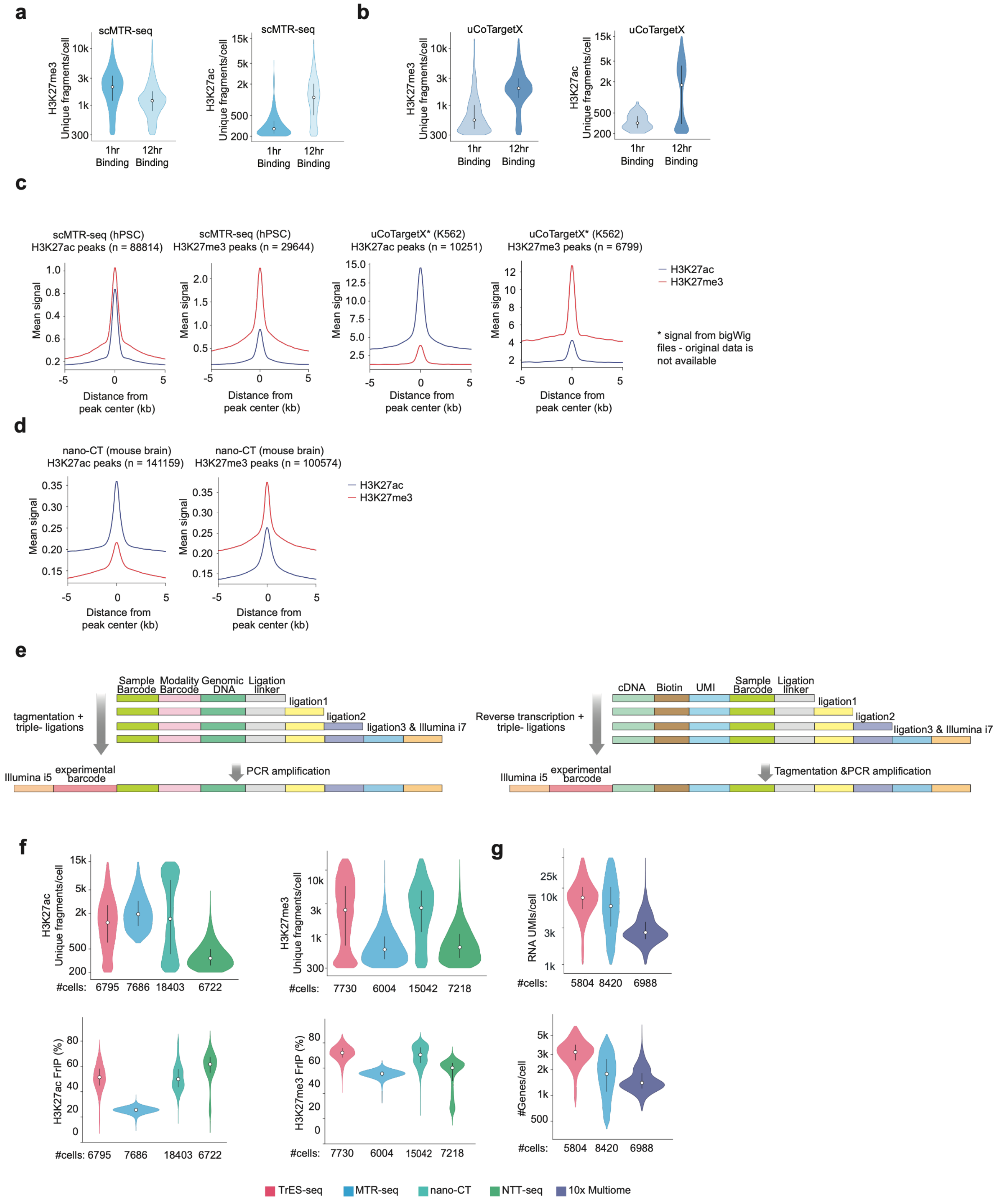

**Suppl. Figure 1**

**a-b.** Violin plots showing number of unique fragment per cell of H3K27me3(left) and H3K27ac(right) in scMTR-seq (a) and uCoTargetX (b) for 1hr and 12hr antibody pATn5 complex binding. **c-d.** Mean H3K27ac and H3K27me3 coverage signal within ±5 kb of peak midpoints for MTR-seq and uCoTargetX (c), nano-CT (d). Profiles are centered on each publicly available dataset's own peaks for the indicated mark. n, number of peaks. **e.** Schematic design of the oligonucleotides used to identify genomic regions marked by dual HPTMs (left) and RNA (right). **f.** Violin plots of single-cell data quality metrics for publicly available datasets from the indicated technologies. Unique fragments per cell (top) and fraction of reads in peaks (bottom) for H3K27ac (left) and H3K27me3 (right). **g.** UMIs (top) and detected genes (bottom) per cell.

Suppl. Figure 2

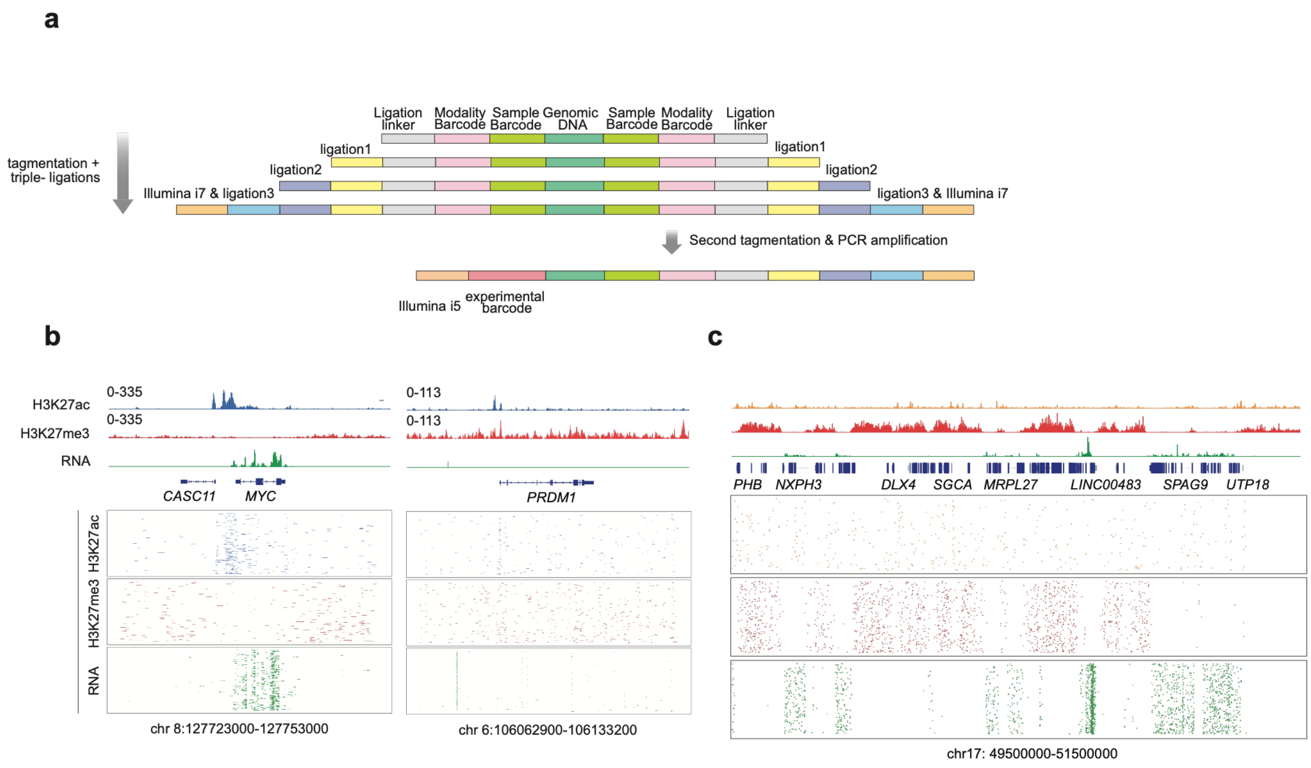

**Suppl. Figure 2**

**a.** Schematic design of the oligonucleotides used for dual tagmentation. **b.** Genome browser views of matched TrES-seq profiles at selected loci in the Karpas-422 cell line. Top aggregated signal tracks for H3k27ac(blue), H3K27me3 (red) and RNA (green) with gene annotations. Bottom: Signal-cell H3K27ac, H3K27me3 and RNA reads across the same regions. Single cell profiles are sorted by descending yield. **c.** Genome browser views of matched TrES-seq profiles at selected loci in the Karpas-422 cell line. Top: aggregated signal tracks for H3K9me3 (orange), H3K27me3 (red), and RNA (green) with gene annotations. Bottom: Single-cell H3K9me3, H3K27me3 and RNA reads across the same regions. Single-cell profiles are sorted by descending yield.

Suppl. Figure 3

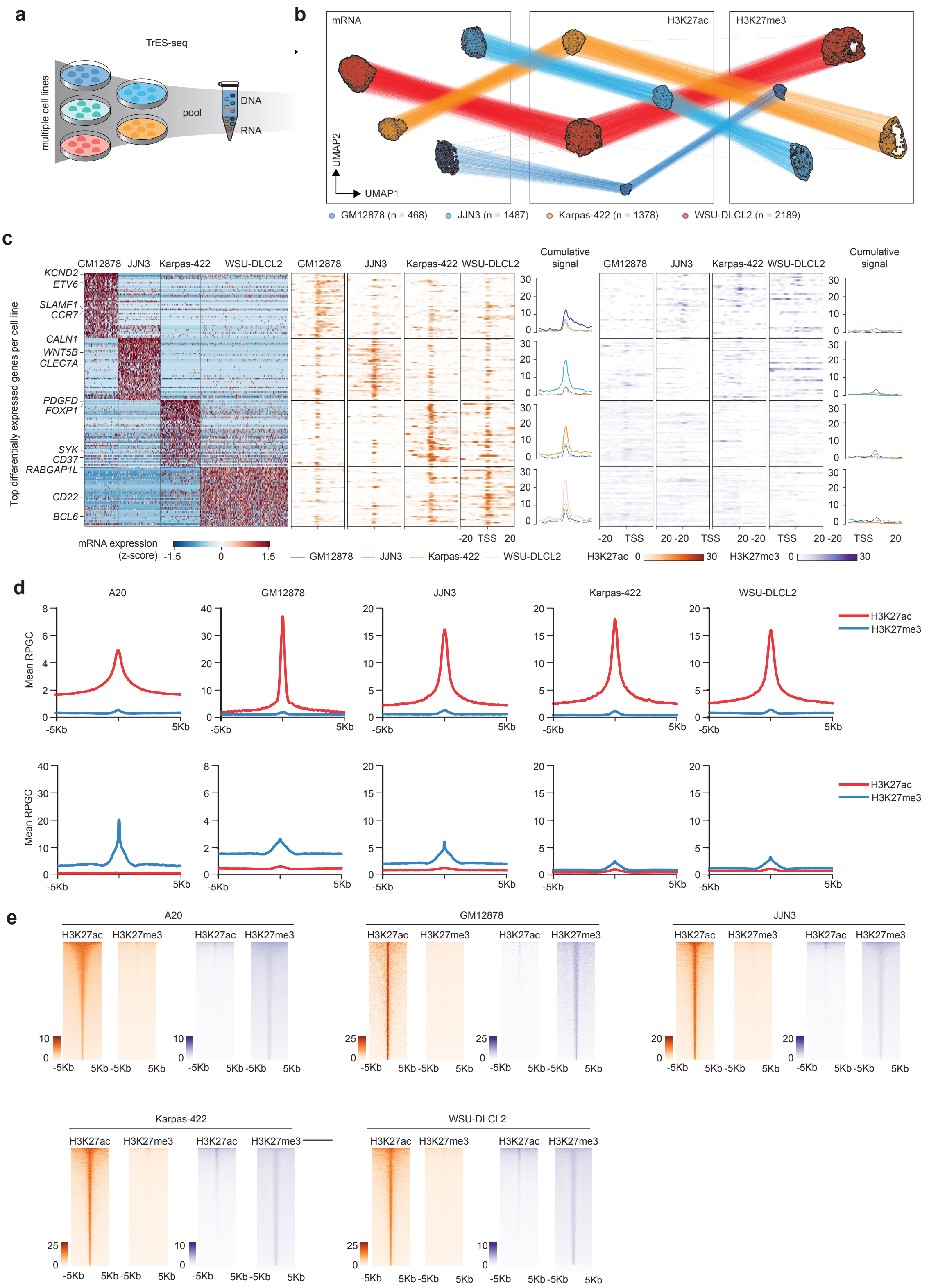

**Suppl. Figure 3**

**a.** Schematic representation of the experimental design used to profile five cell lines: Karpas-422, WSU-DLCL2, GM12878, JJN3, and A20. **b.** UMAP embeddings of the multimodal TrES-seq data for RNA (left), H3K27ac (center), and H3K27me3 (right). Corresponding cells in different modality UMAPs are connected by lines. **c.** Heatmap of the mRNA abundance for the top 50 marker genes of each cell line (left). Total read counts per gene and metagene plots for H3K27ac (center) and H3K27me3 (right) at the transcription start sites (TSS) of the top 50 marker genes per cell line. **d.** Profile plots aggregating the H3K27ac (red) and H3K27me3 (blue) signals at significant H3K27ac (top) and H3K27me3 (bottom) peaks (± 5kb). **e.** For each cell line (indicated in the top), metagene plots showing H3K27ac and H3K27me3 signal intensity at significant H3K27ac (orange) and H3K27me3 (blue) peaks (± 5kb).

Suppl. Figure 4

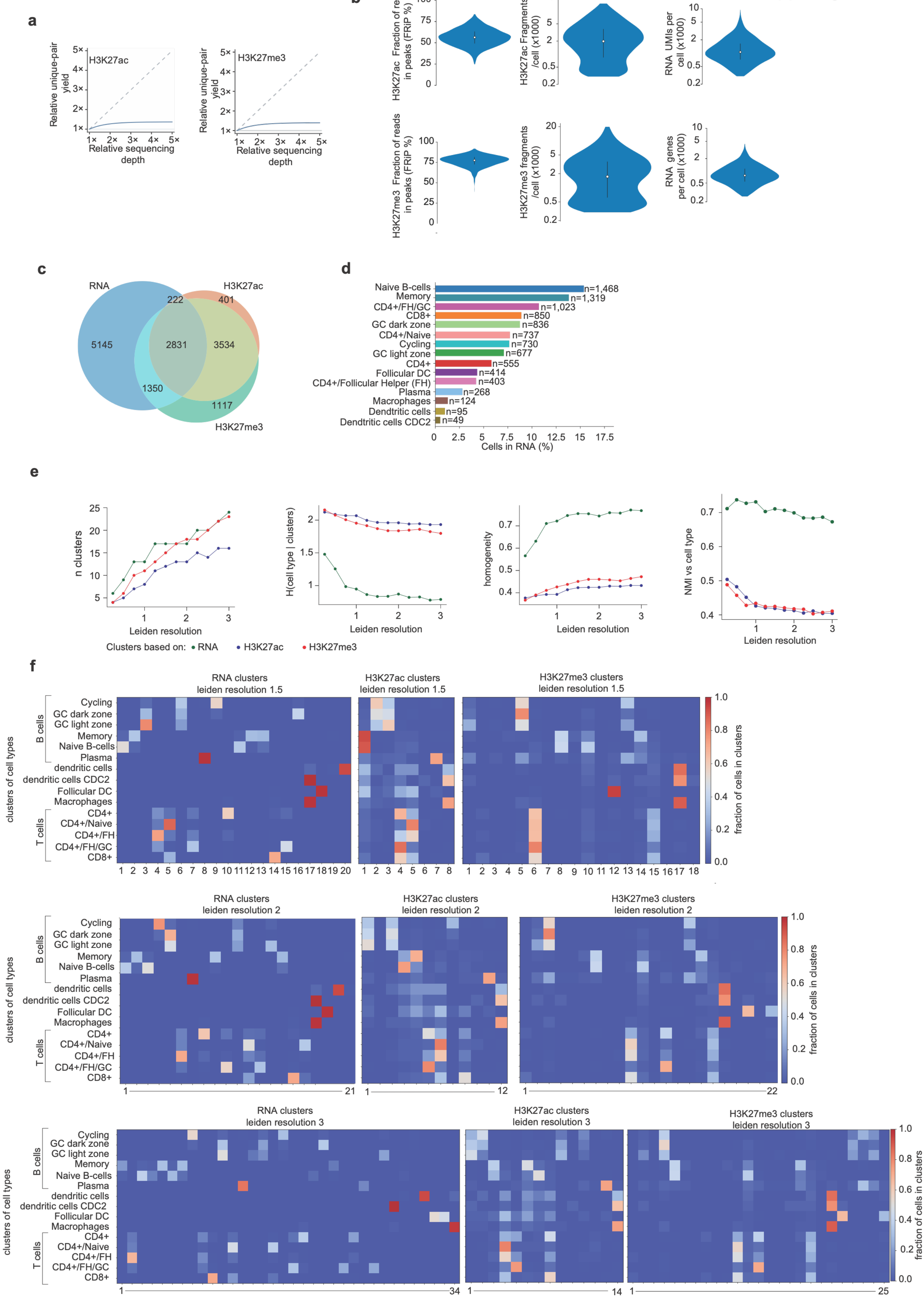

**Suppl. Figure 4**

**a.** Predicted relative yield of unique read pairs with deeper sequencing for H3K27ac (left) and H3K27me3 (right). Solid curves show Picard MarkDuplicates estimates relative to the observed yield at 1× sequencing depth. Dashed lines indicate proportional increases in yield with sequencing depth. **b.** Violin plots showing quality metrics for H3K27ac (top) and H3K27me3 (bottom), FRiP(left), number of unique fragments per cell (middle), and RNA UMIs per cell and genes detected per cell (right). **c.** Venn diagram showing the number of cells passing quality check based on each individual modality and their intersection. **d.** Quantification of the number of cells attributed to each cell subpopulation after label transfer. **e.** Number of clusters (left), and comparison of unbiased clusters vs cell type labels based on: conditional entropy (H), homogeneity, and normalized mutual information (NMI) in the RNA, H3K27ac, and H3K27me3 modalities using the Leiden algorithm at varying resolutions. **f.** Confusion matrices (row normalized) comparing annotated cell types (rows) to unsupervised clusters (columns) for RNA (left), H3K27ac (middle) and H3K27me3 (right) at three clustering resolutions.

Suppl. Figure 5

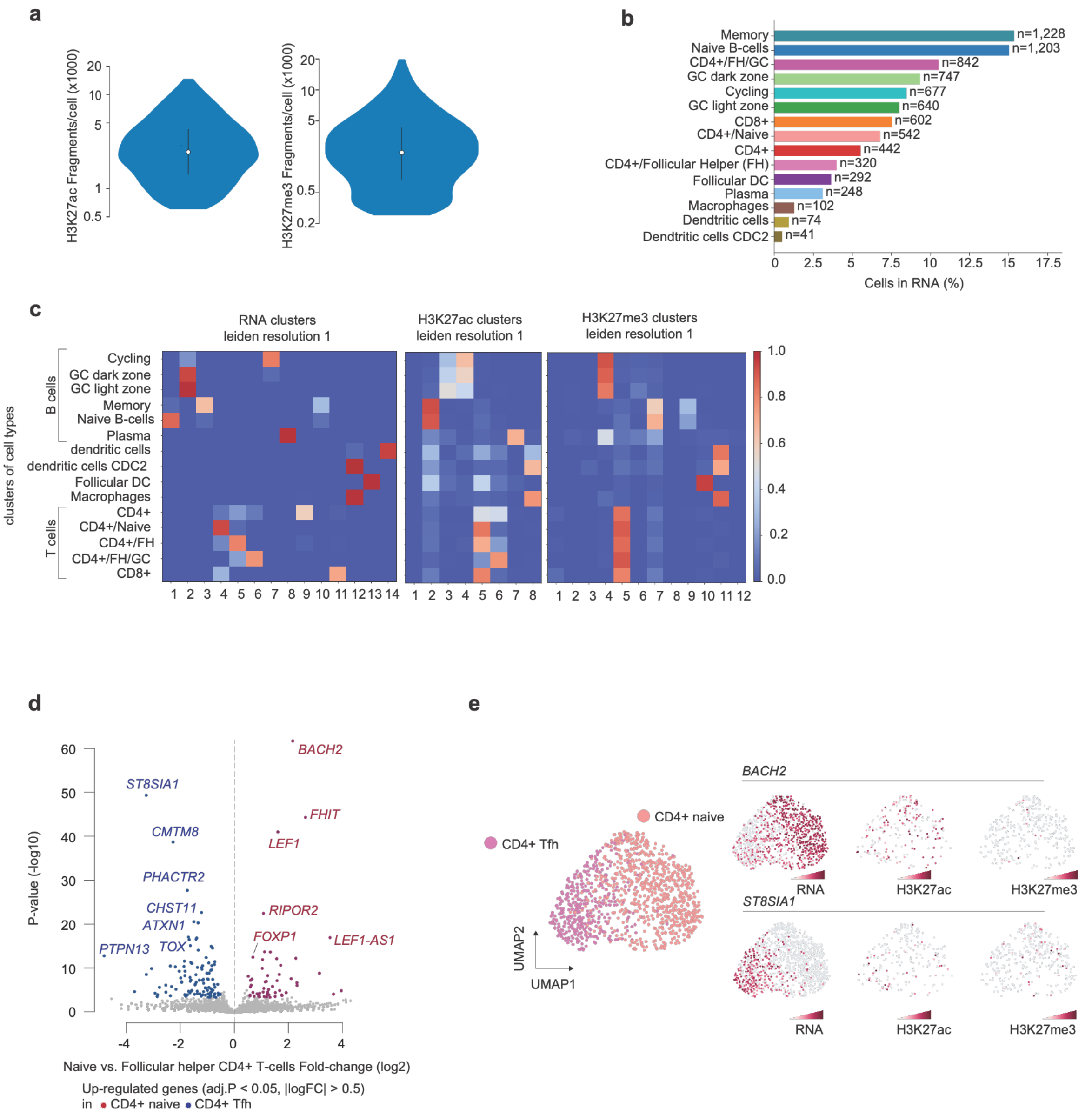

**Suppl. Figure 5**

**a.** Violin plots showing quality metrics of high coverage cells for H3K27ac (left) and H3K27me3 (bottom), number of unique fragments per cell. **b.** Quantification of the number of cells attributed to each cell subpopulation after label transfer. **c.** Confusion matrices (row normalized) comparing annotated cell types (rows) to unsupervised clusters (columns) for RNA (left), H3K27ac (middle) and H3K27me3 (right) at leiden cluster resolution 1. **d.** Volcano plot showing differentially expressed genes between naïve CD4+ and T follicular helper (Tfh) CD4+ cells. **e.** UMAP showing CD4+ naïve and CD4+ Tfh cells showing RNA expression (left), H3K27ac(middle) and H3K27me3(right) signals of BACH2 (top) and ST8SIA1 (bottom). All signals are shown as log1p(CP10K).

Suppl. Figure 6

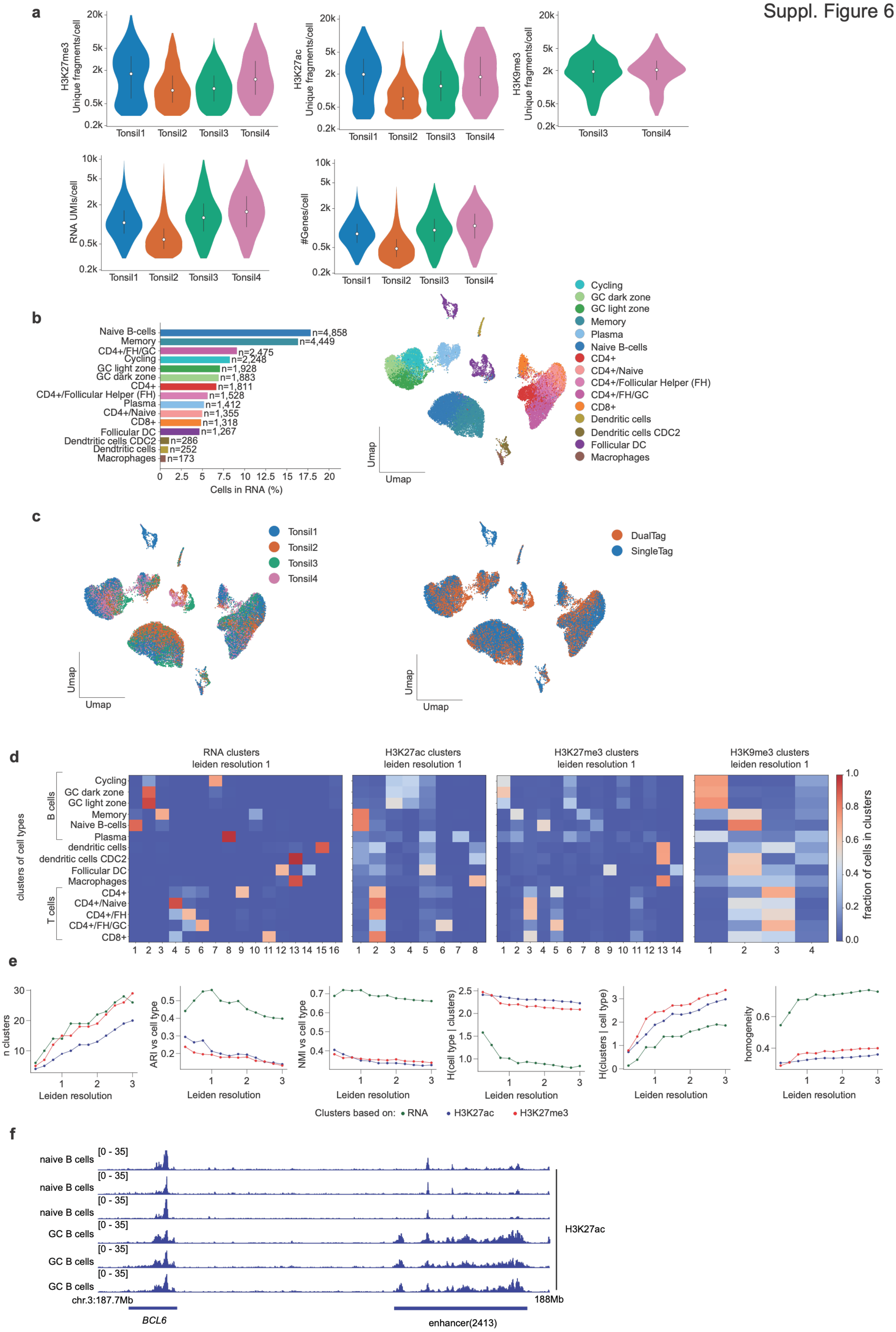

**Suppl. Figure 6**

**a.** Violin plots showing quality metrics across four human tonsil samples, including the number of unique H3K27me3, H3K27ac, and H3K9me3 fragments per cell (top), and RNA UMIs per cell and genes detected per cell (bottom). **b.** Quantification of the number of cells attributed to each cell subpopulation after label transfer. **c.** RNA UMAP of tonsil derived cells colored by donor (left) and TrES-seq single- or dual-tagmentation protocol (right). **d.** Confusion matrices (row normalized) comparing annotated cell types (rows) to unsupervised clusters (columns) for RNA (left), H3K27ac (middle) and H3K27me3 (right) at leiden cluster resolution 1. **e.** Number of clusters (left), and comparison of unbiased clusters vs cell type labels based on: conditional entropy (H), homogeneity, and normalized mutual information (NMI) in the RNA, H3K27ac, and H3K27me3 modalities using the Leiden algorithm at varying resolutions. **f.** Genome browser views of H3K27ac ChIP-seq profiles at the *BCL6* locus and distal enhancer region for the indicated cell types.

Suppl. Figure 7

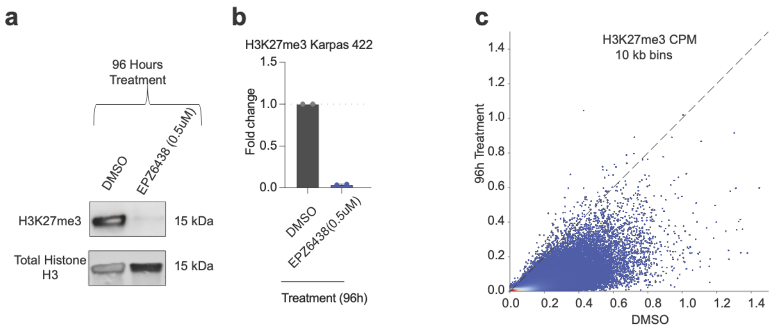

**Suppl. Figure 7**

**a.** Western blot of H3K27me3 and total histone H3 in Karpas-422 cells treated with DMSO or the EZH2 inhibitor EPZ6438 (0.5 μM) for 96 h. **b.** Quantification of H3K27me3 levels normalized to total histone H3, expressed as fold change relative to DMSO-treated cells. **c.** Scatter plot of CPM normalized H3K27me3 signal at 10kb bins in Karpas-422 DMSO (x-axis) vs 96 hours EZH2i treatment (y-axis).
